# *Aspergillus fumigatus* FhdA transcription factor is important for mitochondrial activity and codon usage regulation during the caspofungin paradoxical effect

**DOI:** 10.1101/2022.05.21.492902

**Authors:** Ana Cristina Colabardini, Norman Van Rhijn, Abigail L. LaBella, Clara Valero, Fang Wang, Lauren Dineen, Michael J. Bromley, Antonis Rokas, Gustavo H. Goldman

## Abstract

*Aspergillus fumigatus* is the main etiological agent of aspergillosis. The antifungal drug caspofungin (CSP) can be used against *A. fumigatus* and CSP tolerance is observed. Here, we show that the transcription factor FhdA is important for mitochondrial activity and regulates genes transcribed by RNA polymerase II and III. FhdA influences the expression of tRNAs that are important for mitochondrial function upon CSP. Our results show a completely novel mechanism that is impacted by CSP.

## INTRODUCTION

*Aspergillus fumigatus* is a filamentous saprophytic fungus and an opportunistic pathogen (1). It is the main pathogen responsible for invasive pulmonary aspergillosis (IPA), one of the most severe infections in immunosuppressed patients in terms of morbidity and mortality (2). The echinocandin caspofungin (CSP) is a fungistatic drug against filamentous fungi and can be used as salvage therapy for IPA (3). It acts by non-competitively inhibiting the fungal β-1,3-glucan synthase (Fks1), which is required for the biosynthesis of the primary fungal cell wall carbohydrate β-1,3-glucan (4). However, in a certain range of higher concentrations, there is a reduction of CSP activity. This phenomenon, which is known as “caspofungin paradoxical effect” (CPE) and results from a tolerance cellular response that alters the cell wall content and fungal growth (5).

*A. fumigatus* is a highly successful opportunistic pathogen mostly due to its ability to rapidly adapt to diverse environments. To this end, changes in physicochemical conditions and nutrient availability generate signals at the cell surface that are conveyed by a system of signaling pathways to the nucleus and converge at transcription factors (TFs) (1). TFs regulate the transcription of gene sets that drive metabolic reprogramming to enable adaptation to the new conditions and promote proliferation inside the host (6). Recently, by screening a library of 484 TF null mutants (7), we identified FhdA (AFUB_091020), a novel TF that plays a role in CSP response (8). The observations that the Δ*fhdA* strain is more sensitive to CSP and lacks the CPE (Figure 1A), and that FhdA is constitutively located inside the nucleus (Figure 1B), led us to further function characterization of this TF.

**Figure 1:**
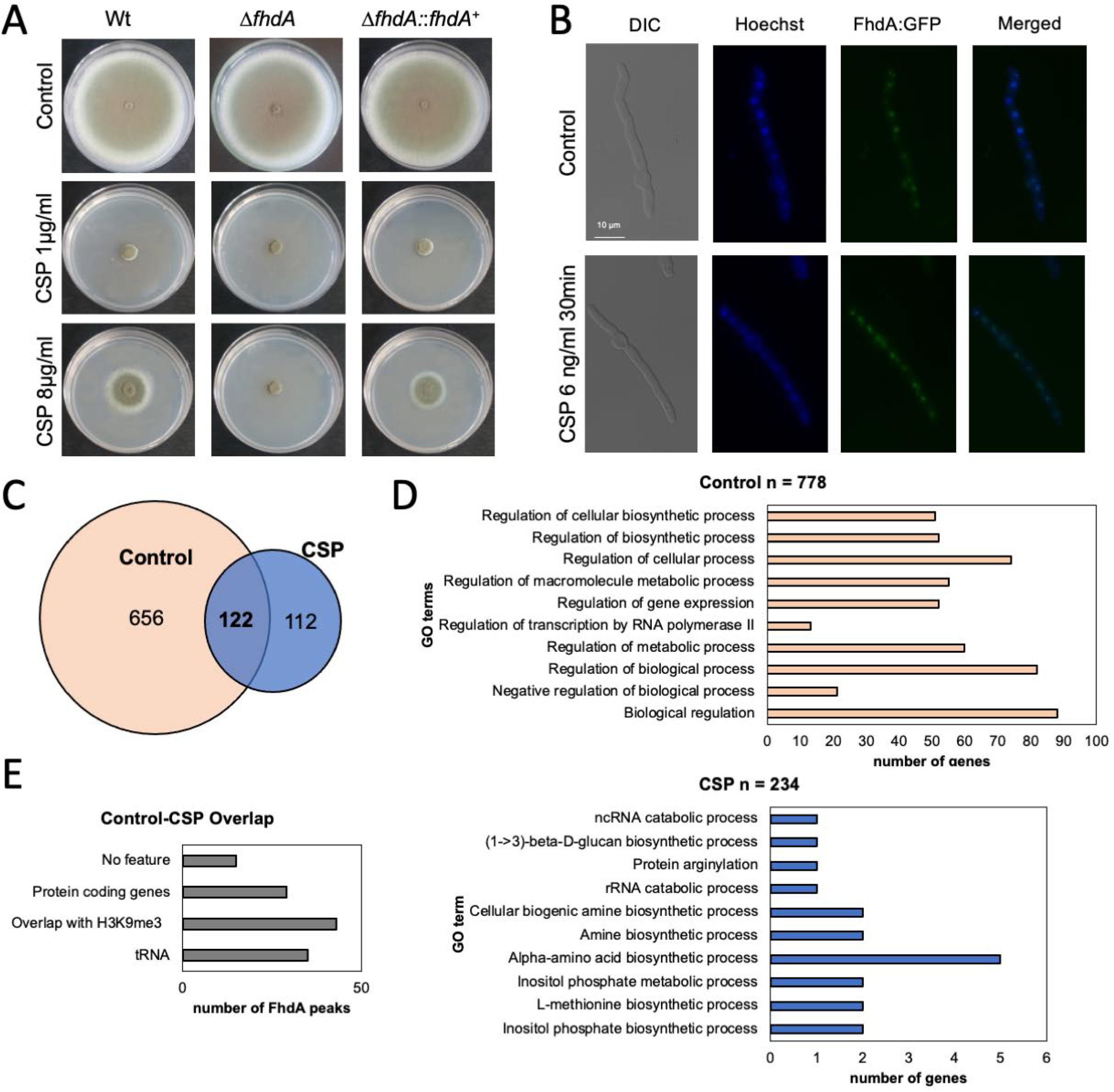
FhdA is essential to the caspofungin paradoxical effect (CPE) and binds to several targets before and after caspofungin (CSP) exposure. (A) Radial growth of the wild-type strain (Wt, CEA17), Δ*fhdA*, and Δ*fhdA:fhdA*^+^ incubated for 5 days at 37°C in minimal medium containing 0, 1 and 8 µg/ml of CSP. (B) Fluorescence microscopy of a FdhA:GFP functional strain before and after exposure to 6ng/ml of CSP for 30 minutes. Hoescht was used to dye the nuclei. (C) A Venn diagram showing the number of FhdA targets before and after exposure to 2 µg/ml of CSP for 1h. Heatmaps of ChIP-seq peak intensity of FhdA to *A. fumigatus* tRNA genes in control and caspofungin conditions. The average peak intensity was taken for two replicates and 1kb up and downstream of the centre of each tRNA gene was used for visualisation using DeepTools. Higher peak intensities for tRNAs in caspofungin compared to control conditions. In general, binding of FhdA was found within the tRNA coding region. However, several peaks upstream of tRNA genes could be identified. The methodology for the ChIP-seq is described in Colabardini *et al*. (2022). (D) Top ten GO terms categorizing the genes closest to the FhdA targets before and after CSP exposure. (E) Functional classification of the 122 targets constitutively bound by FhdA.

In order to explore the pathways affected by FhdA, we investigated its direct targets by determining the FhdA:3xHA binding sites using genome-wide ChIP-seq (chromatin immunoprecipitation coupled to DNA sequencing) analysis. We detected a total of 890 regions bound by FhdA at their promoters (Figure 1C). Most of the promoters (n=778) were bound by FhdA before CSP exposure, of which 122 were constitutively bound by the TF both before and after exposure to 2 µg/ml of CSP (Figure 1C). In contrast, 112 genes were bound by FhdA exclusively after CSP exposure (Figure 1C).

Enrichment analysis of the 778 genes bound by FhdA in control conditions suggest that this TF is involved in regulation of processes related to gene expression and transcriptional control (Figure 1D, upper panel), while the 234 genes bound after CSP exposure are mostly involved in amino acid metabolism (Figure 1D, bottom panel). Of the 122 constitutive targets of FhdA, 15 have no feature, 29 encode proteins, 43 overlap with intergenic regions bound by the modified histone H3K9me3 previously described by our group (9), and 35 encode tRNAs (Figure 1E).

FhdA is important for mitochondrial respiratory function (Valero *et al*., 2020). We analyzed the 29 protein coding genes bound by TF using MitoProt (https://openebench.bsc.es/tool/mitoprot_ii) revealing eight that had a mitochondrial targeting signature. These are a putative 30S ribosomal subunit S4 (AFUB_007360), the TF LeuB (AFUB_020530), the putative ER Hsp70 chaperone BipA (AFUB_021670), the putative ABC transporter Adp1 (AFUB_073240), a ketol-acid reductoisomerase (AFUB_034740), a putative glucosamine-6-phosphate deaminase (AFUB_083490), a protein predicted to bind chromatin (AFUB_096570), and a hypothetical protein (AFUB_077560). It remains to be determined if these proteins localize at the mitochondria and how they affect CSP tolerance.

We then hypothesized that the tRNA genes controlled by FhdA could influence the mitochondrial tRNA pool. *A. fumigatus* has 27 predicted mitochondrially-encoded tRNA anti-codons with 47 codons that can be decoded from wobble or exact match pairing. That leaves 13 codons that cannot be decoded by the mitochondrially-encoded tRNAs without tRNA modifications (Figure 2A). Seven of these 13 codons without a tRNA encoded in the mitochondria can be decoded by one of nine tRNAs identified in the ChIP-seq experiment (Figure 2B).

**Figure 2:**
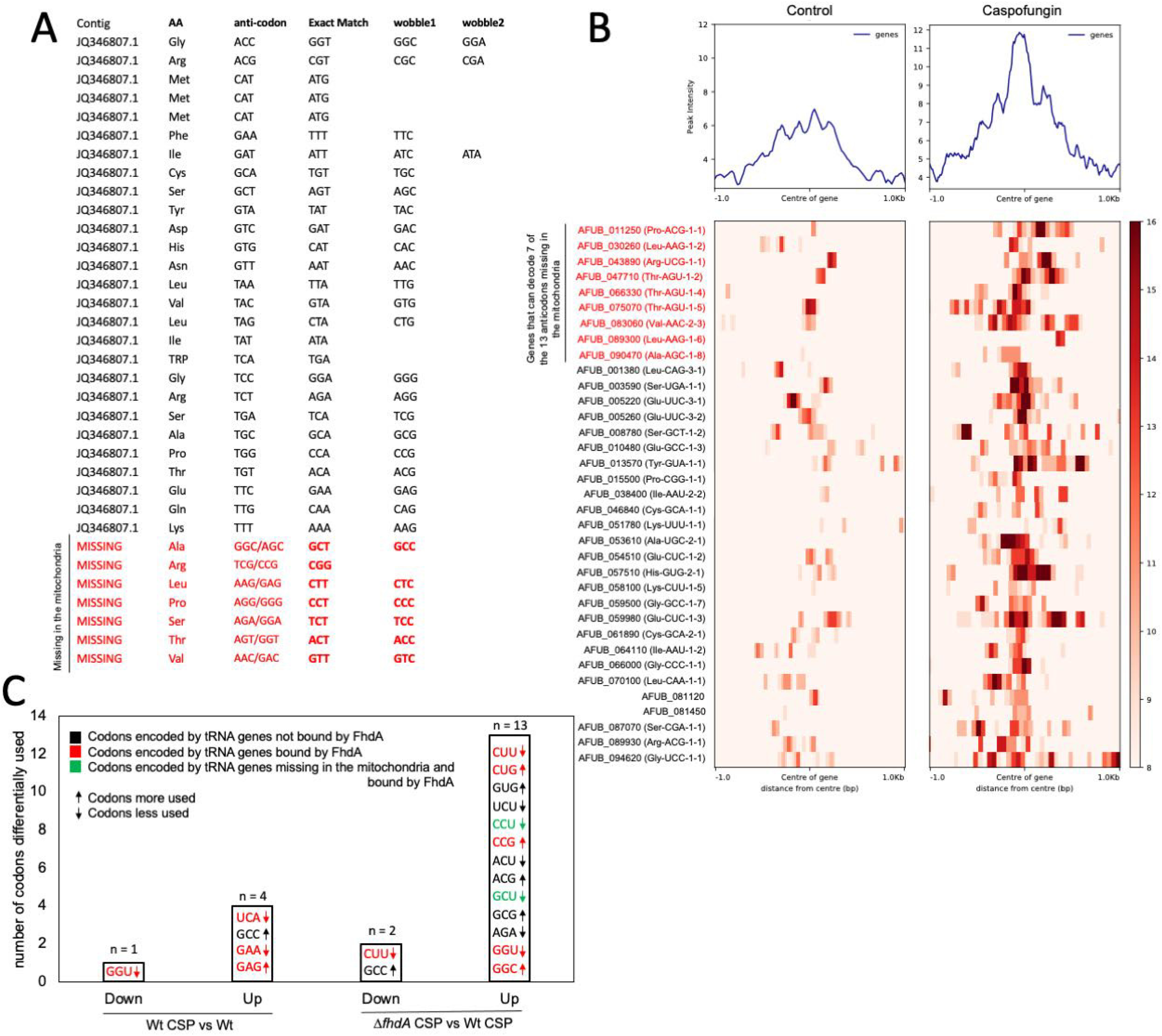
FhdA binds tRNA promoters, including promoters of several tRNAs that are not encoded by the mitochondrial genome. (A) The tRNA genes encoded by the mitochondrial genomes and the ones that are absent from it. (B) A heat map representing the FhdA binding intensity to the 35 tRNA genes detected in our ChIP-seq. (C) Relative synonymous codon usage (RSCU) analysis showing the number of codons differentially used for the transcription of the 5-fold up- and down-regulated genes detected in the RNA-seq experiment.

We then asked if the *fhdA* deletion effect could be related to an altered tRNA preference. To test for codon specific effects on nuclear gene expression we compared the Relative-Synonymous-Codon-Usage (RSCU), defined as the ratio of the observed frequency of codons to the expected frequency given that all the synonymous codons for the same amino acids are used equally (10), using the RNAseq dataset where genes regulated by FhdA in the absence or presence of CSP are 5-fold up or down regulated (8). When we compared the wild-type exposed to CSP with the corresponding control, genes that were 5-fold up-or down-regulated during exposure to CSP had lower usage of the codons UCA, GAA, and GGU and higher usage of the codons GAG and GCC. Four of these five codons are decoded by tRNAs identified in the ChIP-seq experiment (Figure 2C). Genes over or under-expressed in the Δ*fhdA* mutant when compared to the wild-type, both treated with CSP, were enriched or depleted for 15 different codons in up or down-expressed genes; eight of these codons are decoded by tRNAs identified in the ChIP-seq and two of these codons are not present in the mitochondria (Figure 2C). These results suggest that there are differences in codon usage during CSP response; there are about 3-fold more differences in codon usage in Δ*fdhA* during CSP, suggesting that the combination of Δ*fhdA* and CSP affects gene expression in a codon-specific manner that may be related to changes in tRNA expression.

In summary, our results suggest that CSP can modulate the tRNA pool utilization, indicating that several side effects related to CSP activity, such as a decrease in the mitochondrial function could be caused by depletion of essential tRNAs. FhdA seems to play an important role in the control of the expression of protein encoded genes but also tRNA expression, suggesting that FhdA can influence RNA PolII and RNA PolIII targets. Transcriptional regulation of tRNAs may represent a novel route of translation regulation in filamentous fungi.

## ACKNOWLEDGEMENTS

We thank the Fundação de Amparo à Pesquisa do Estado de São Paulo (FAPESP) grants numbers 2017/07536-4 (A.C.C.), 2018/00715-3 (C.V.), and 2016/07870-9 (G.H.G.) and the Conselho Nacional de Desenvolvimento Científico e Tecnológico (CNPq) grant numbers 301058/2019-9 and 404735/2018-5 (G.H.G.), both from Brazil, and the National Institutes of Health/National Institute of Allergy and Infectious Diseases (R01AI153356) from the USA (A.R. and G.H.G.). This work was also supported by the Wellcome Trust grants no. 219551/Z/19/Z and 208396/Z/17/Z to M.B.

